# Site-specific C-terminal fluorescent labeling of Tau protein

**DOI:** 10.1101/2022.06.07.495075

**Authors:** Louise Bryan, Saurabh Awasthi, Yuanjie Li, Peter Niraj Nirmalraj, Sandor Balog, Jerry Yang, Michael Mayer

## Abstract

Formation of Tau protein aggregates in neurons is a pathological hallmark of several neurodegenerative diseases, including Alzheimer’s disease. Fluorescently-labeled Tau protein is therefore useful to study the aggregation of these pathological proteins and to identify potential therapeutic targets. Conventionally, cysteine residues are used for labeling Tau proteins, however, the full-length Tau isoform contains two cysteine residues in the microtubule-binding region, which are implicated in Tau aggregation by forming intermolecular disulfide bonds. In order to prevent the fluorescent label from disturbing the microtubule binding region, we developed a strategy to fluorescently label Tau at its C-terminus while leaving cysteine residues unperturbed. We took advantage of a Sortase A-mediated transpeptidation approach to bind a short peptide (GGGH6-Alexa_647_) with a His-tag and a covalently attached Alexa 647 fluorophore to the C-terminus of Tau. This reaction relies on the presence of a Sortase recognition motif (LPXTG), which we attached to the C-terminus of recombinantly expressed Tau. We determined the possible effects of the resulting C-terminal modification on the secondary structure of Tau protein, its aggregation kinetics, and fibril morphology compared to the unlabeled native Tau protein *in vitro*. The results showed no significant differences between the native and C-terminally labeled Tau monomer with regard to aggregation kinetics, secondary structure, and morphology.

## Introduction

Tauopathies are a group of diseases affecting the brain caused by the aggregation of microtubule-associated protein Tau.^1, 2^ Alzheimer’s disease (AD) is one of the most common neurodegenerative disorders, characterized by progressive memory loss and cognitive dysfunction and Tau proteins are increasingly implicated in AD as well as other neurodegenerative diseases.^3, 4^ Tau proteins are predominantly found in neuronal cells and are essential for the assembly and stabilization of microtubules.^5, 6^ Human Tau protein exists in six different isoforms ranging in length (from 352 to 441 amino acids) that vary in the number of N-terminal inserts (N1, N2) and contain 3 or 4 imperfect repeats (R1-R4). Repeats R1-R4 correspond to the microtubule-binding region (**Figure 1A**).^7, 8^ The soluble monomeric form of Tau is thought to be intrinsically disordered and can exist in different conformations.^9, 10^ Numerous post-translational modifications of Tau protein, such as phosphorylation, acetylation, or methylation, regulate its interactions with microtubules or other proteins.^11, 12^ Phosphorylation is one of the most prevalent modifications with up to 85 potential sites on serine, threonine, or tyrosine residues.^13^ Hyper-phosphorylation of Tau results in impaired microtubule binding and leads to misfolding and aggregation, affecting neuronal stability and causing neuronal loss.^14, 15^ The exact aggregation mechanism of Tau and hyper-phosphorylated Tau is still under investigation; however, it is thought to undergo a nucleation-dependent pathway.^16^ Studies have suggested that Tau also undergoes liquid-liquid phase separation, a property of intrinsically disordered proteins (IDPs).^17^ The most toxic species are early aggregates of Tau protein, i.e., soluble or low-n oligomers.^18, 19^ These small aggregates self-assemble into fibrils and finally into neurofibrillary tangles.^2^ High-resolution cryo-electron microscopy (cryo-EM) images of purified Tau aggregates from the brains of AD patients display an ordered core of pairs of protofilaments comprising regions R3 and R4 (**Figure 1B**), with the disordered N- and C-termini forming so called fuzzy coat.^20,21^

**Figure 1.**
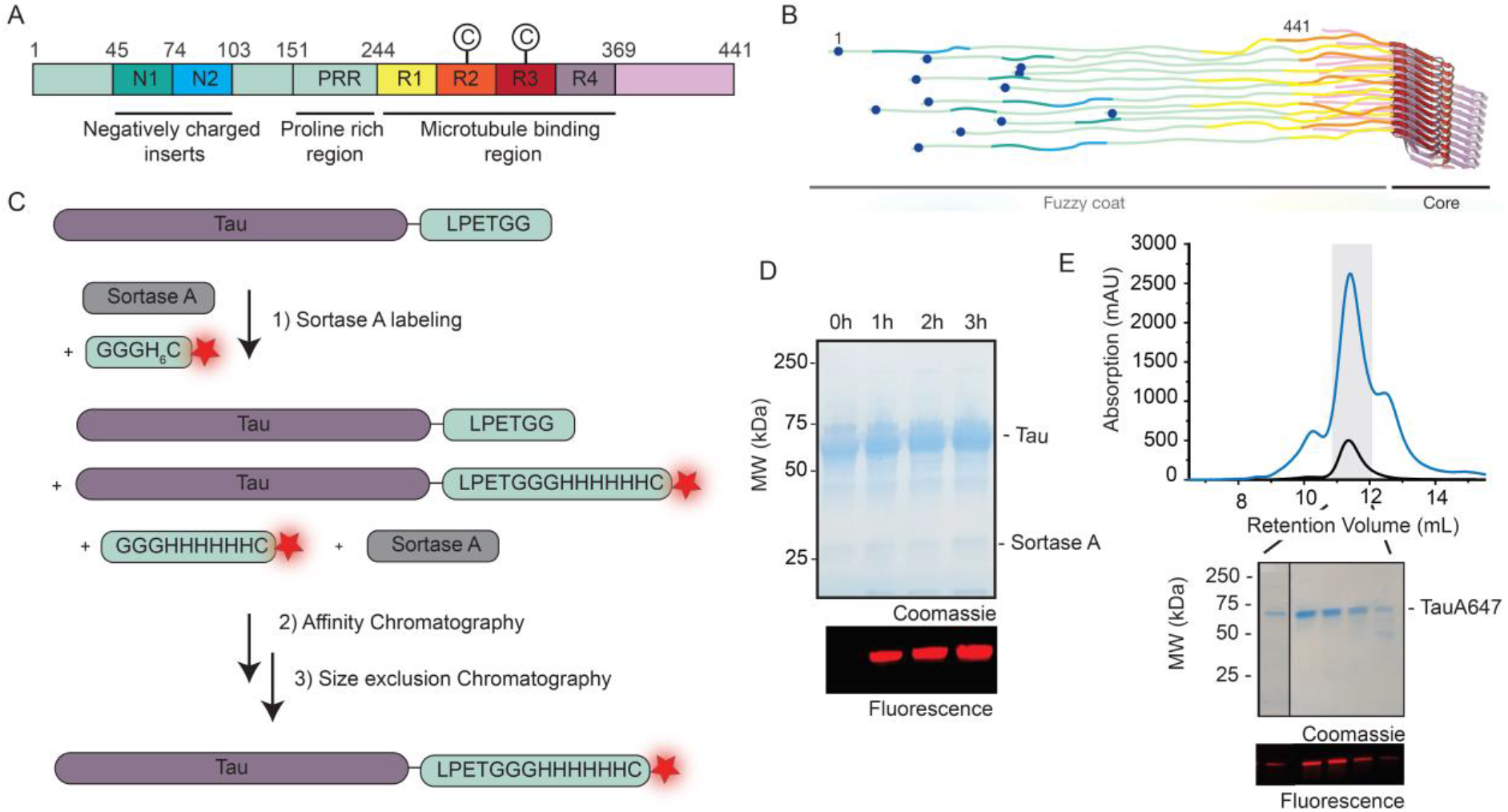
Full length Tau protein and strategy for C-terminal labeling. **A)** Schematic representation of full-length Tau protein (2N4R) colored by domain. N1, N2: N-terminal inserts, PRR: proline rich region, R1-4: imperfect regions,©indicates cysteine residues **B)** Schematic structure of tau fibril consisting of a core and fuzzy coat. Reprinted by permission from Macmillan Publishers Ltd: copyright 2017.^22^ **C)** Schematic representation of C-terminal labeling of Tau protein with the enzyme Sortase A. The recombinantly expressed full-length Tau protein is extended on its C-terminus by addition of an -LPETGG peptide. Sortase A catalyzes the attachment of a his-tag with a covalently attached Alexa647 fluorophore. **D)** Top: Coomassie blue staining of an SDS-PAGE gel of Tau protein after a reaction at time 0, 1h, 2h and 3h upon addition of Sortase A and labelled peptide; bottom: fluorescence image of the Tau band. **E)** Size exclusion chromatography trace (FPLC) of Tau-Alexa647 purification (blue: absorbance at 214 nm, black: absorbance at 647 nm) and SDS-PAGE gel with top: coomassie stain and bottom: fluorescence of Tau-Alexa647.

Single-molecule fluorescence microscopy has the potential to improve our understanding of Tau aggregation. Labeling of Tau protein with a high-quality fluorophore such as the Alexa647 dye used here is essential for such studies. It is, however, important to consider two key factors when designing a method to label Tau protein: First, the presence of two hydrophobic hexapeptide sequences; _275_VQIINK_280_ (known as PHF6*) in R2 and especially _306_VQIVYK_311_ (PHF6) in R3,^23^ which are strongly associated with Tau protein aggregation and should not be perturbed.^24, 25^ Second, the presence of two cysteine residues (at position 291 and 322) (**Figure 1A**), which are involved in intermolecular disulfide bond formation should be preserved. In particular Cys-322 is considered to drive the initial dimerization of Tau protein monomers and aggregation into paired helical filaments.^26-28^ Since modification of these cysteine residues with a fluorophore may influence the aggregation mechanism of Tau protein, we sought a strategy to label the protein on either the N- or C-terminus. Both termini are part of the fuzzy coat and are located away from the core region of Tau fibrils (**Figure 1B**). To our knowledge, labeled Tau has so far only been prepared using K_18_ or shorter Tau isoforms by modification of amine groups or by site-specific mutagenesis of one of the two cysteine residues in the core region of full-length Tau.^29-33^

Here, we employed a site-specific approach to label the C-terminal end of full-length Tau protein and investigated the effects of modifications on its secondary structure and aggregation kinetics compared to those of native Tau protein *in vitro*. We monitored the aggregation of native and fluorescently labeled Tau over time in the presence of polyanionic inducer heparin^34^ using a ThT assay.^35^ In addition, we compared the morphology of the resulting Tau fibrils using transmission electron microscopy (TEM) and atomic force microscopy (AFM).^36-39^ We observed that the native and C-terminally labeled Tau protein exhibit similar properties with respect to secondary structure, fibril morphology and aggregation kinetics based on analysis by CD Spectroscopy and imaging by TEM and AFM.

## Results and Discussion

### C-terminal labeling of Tau protein

We explored a Sortase A-mediated labeling strategy to fluorescently label Tau protein while causing minimal perturbation at the core region of the protein in the fibrillar state.^40-42^ The method relies on the addition of a specific recognition motif LPXTG (X being any amino acid) to the C terminus of recombinantly expressed Tau protein. Sortase A from *Staphylococcus aureus* cleaves off the terminal glycine and forms a thioester with threonine. This step is followed by conjugation to a (G)_n_ peptide/protein. We thereby used full-length Tau with the addition of the short sequence LPETGG at its C-terminus and enzymatically ligated it to a short peptide bearing a His-tag and a covalently attached fluorescent molecule: GGGH_6_C-Alexa647 (**Figure 1C**). We chose Alexa647 since it is a high-quality, photostable, far-red fluorescent dye and to prevent spectral overlap with Thioflavin T whose fluorescence can be excited from 385 nm to 450 nm and whose emission maximum ranges from 445 nm to 482 nm.^43^ We added the hexahistidine sequence to the peptide to provide an additional tag for efficient separation of the labeled from the unlabelled protein and to concentrate labeled Tau species since the Sortase A labeling reaction typically provides labeling efficiencies of 20 to 90%; largely dependent on the Sortase A variant.^44^ We used the commercially available Sortase pentamutant (Sortase 5A, with a His-tag), which increases the rate and efficiency of Sortase labeling compared to the wild type Sortase A.^45^ The labeling reaction was carried out at 10°C to prevent degradation of the Tau protein or the formation of small aggregates. The reaction mixture was then purified on a Ni-NTA column where the labeled protein, GGGH_6_-Alexa647 and Sortase A, all containing His-tags were retained on the column. This affinity chromatography was followed by size exclusion chromatography to purify the labeled protein from the remaining molecules in the mixture (**Figure 1D and 1E**). This procedure purified C-terminal labeled Tau protein for further structural characterization of the monomeric form of Tau and its aggregates.

### Secondary structure and aggregation kinetics of labeled and native full-length Tau

We determined the secondary structure of monomeric Tau-A647 and native full-length tau protein by CD spectroscopy. **Figure 2A** shows a minimum at 200 nm for both native Tau and Tau-A647, implicating that they are unstructured in solution and the addition of a short peptide sequence with a fluorescent dye on the C-terminus did not change the secondary structure. Furthermore, we analyzed the CD spectra and determine the propensity of different secondary structural elements such as α-helix, β-sheet, and irregular or disordered state for native and labeled Tau protein monomers. Table 1 reveals a similar secondary structure in both the variants (i.e. native and labeled), suggesting a minimal effect of C-terminal labelling on Tau protein structure in solution (**Figure 2A**).

**Table. 1.** Fraction of different secondary structures for native and labeled Tau protein.

| Protein variant | α-Helix (%) | β-Sheet (%) | Irregular (%) |
| --- | --- | --- | --- |
| Native Tau | 3.4 ± 2.5 | 44.0 ± 3.6 | 60.0 ± 3.6 |
| Labeled Tau | 4.7 ± 4.2 | 42.7 ± 7.3 | 57.0 ± 2.6 |

**Figure 2.**
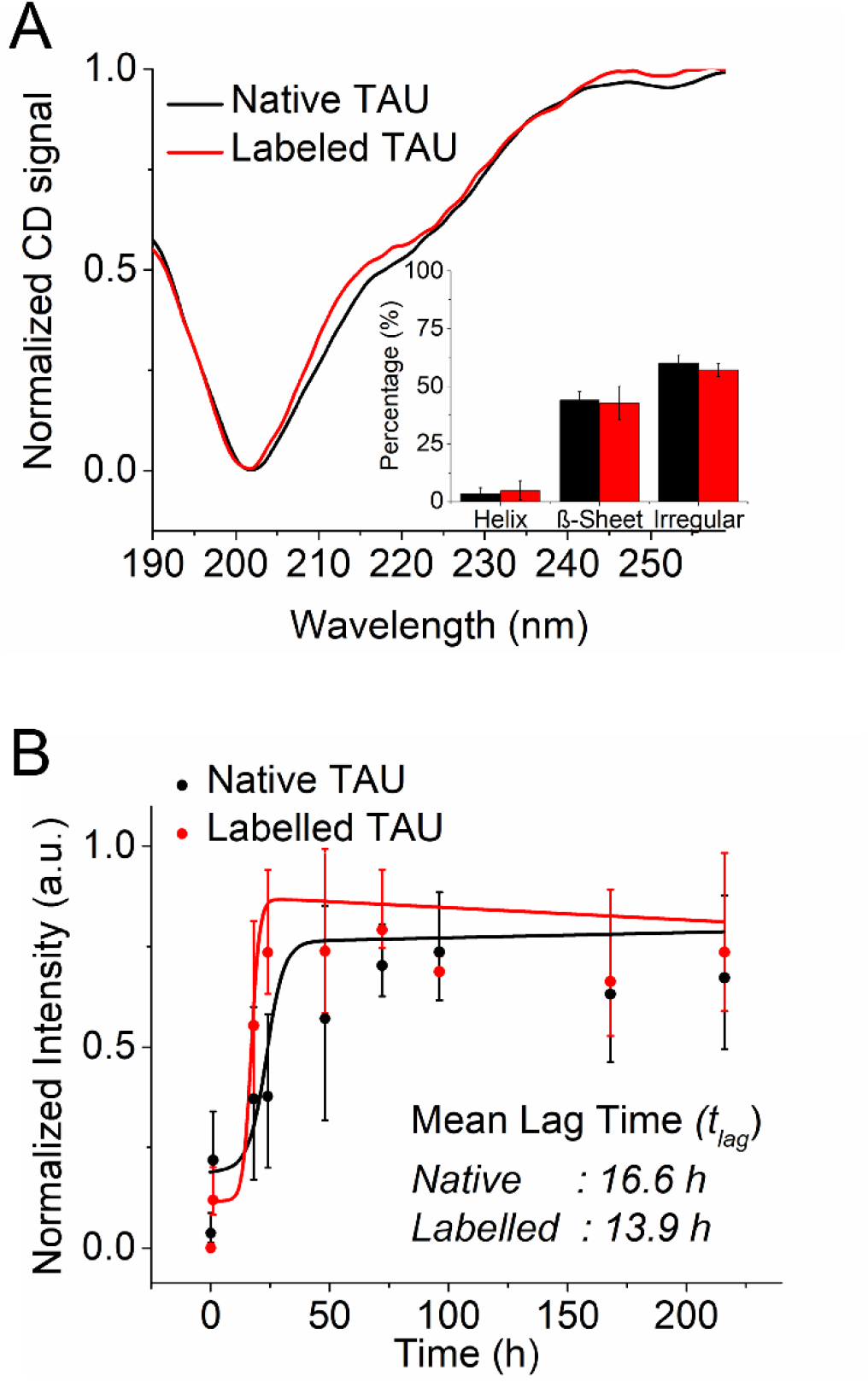
Comparison of the monomer secondary structure of native and labeled Tau in solution as well as their aggregation kinetics. **A)** CD spectra of native (black), and labelled Tau (red) in their predominantly monomeric form. The inset shows the percentage of different secondary structures for native and labeled Tau as determined using the online server CAPITO.^46^ **B)** ThT fluorescence intensity measurements to compare the aggregation kinetics of native and labeled Tau protein. The data points are the average of three independent measurements. The error bars show the standard deviation from three independent measurements.

We also carried out a comparative analysis of aggregation kinetics of labeled and native full-length Tau. In order to follow their aggregation kinetics, each Tau variant was incubated with the inducer, i.e., heparin (a Tau:heparin ratio of 4:1) and ThT at 37°C in Tau aggregation buffer and their fluorescence spectrum was measured at regular time points (**Figure 2B**). Overall, we observed a similar trend in the case of native and labelled Tau protein with the ThT fluorescence increasing overtime corresponding to the formation of aggregated species. We estimate the lag time (*t*_lag_) of aggregation using Equation 1:

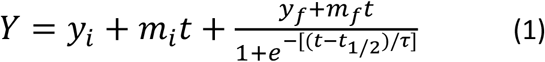

where, *Y* is the fluorescence intensity as a function of time *t, y*_*i*_ and *y*_*f*_ are the intercept of the initial and final fluorescence values with the y-axis, *m*_*i*_ and *m*_*f*_ are slopes of the initial and final baselines, *t*_½_ is the time needed to reach halfway through the elongation phase and τ is the elongation time constant. The apparent rate constant, *k*_app_, for the growth of fibrils is given by 1/s, and the lag time is usually defined as *t*_*lag*_ = *t*_½_ - 2τ.^47^ This analysis revealed that *t*_lag_ was 16.6 h for native Tau and 13.9 h for labeled Tau.

### Labeled and native Tau exhibit similar fibril morphology

In order to compare the fibril morphology, we carried out TEM and AFM imaging of aggregates formed by native and labeled full-length Tau protein. **Figure 3** shows that TEM imaging of Tau fibrils after 7 and 14 days of aggregation *in vitro* induced by heparin revealed similar fibril morphology. From these images, we determined the width of fibrillar aggregates. Both the native full length-Tau protein and labeled Tau-A647 exhibit a similar fibril width estimates of 14.55 ± 1.8 nm and 14.02 ± 1.7 nm, respectively (**Figure 3C and 3F**). Earlier studies have shown that Tau fibrils extracted from brain samples exhibit a width of 19 nm, whereas in the case of heparin-induced Tau protein fibrils they were around 16.9 nm.^48^ More recently, Zhang *et al* showed heparin induced Tau fibrils are in fact highly heterogeneous with a width ranging from 4 to 25 nm.^49^ Our estimates of fibrillar width for both the native and labeled Tau protein are in good agreement with these earlier reports.

**Figure 3.**
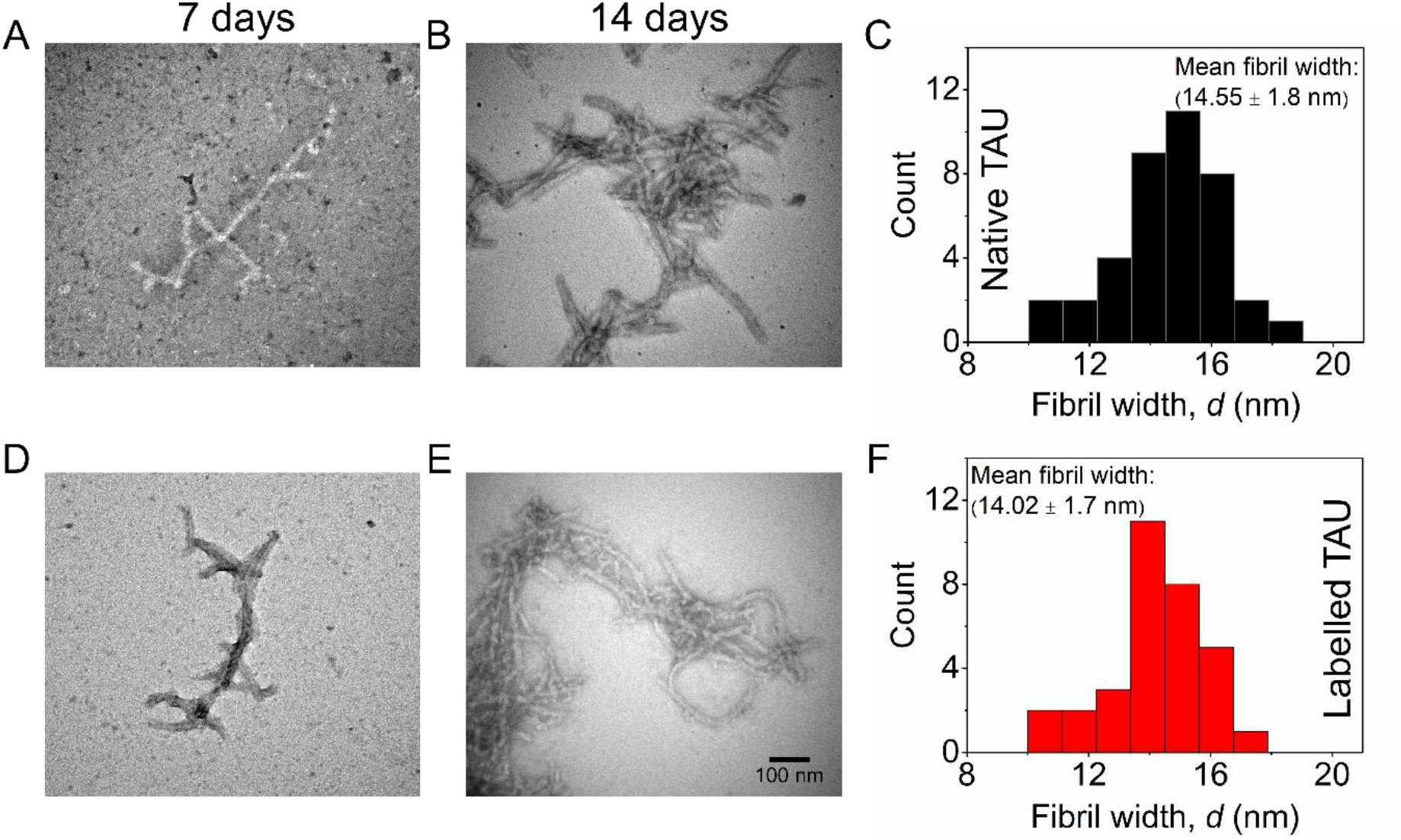
Transmission electron microscopy (TEM) imaging and characterization of fibril width. **A)** and **B)** TEM images of native Tau protein aggregates incubated for 7 days and 14 days at 37°C under shaking conditions, in the presence heparin. **D)** and **E)** TEM images of labelled Tau protein incubated for 7 days and 14 days respectively. (Scale bar: 100 nm). **C)** and **F)** Histograms of the fibril width after 14 days of aggregation.

Furthermore, we characterized the fibrils formed after 7 days of incubation by labeled and native Tau protein using liquid-based AFM to estimate their height (**Figure 4 A-F**). Histograms of the height of individual fibrils revealed a height range of 5.0 ± 1.1 nm for native Tau and 5.6 ± 1.2 nm for Tau-A647 (**Figure 4F**). In general, the high-resolution AFM images do not show any drastic differences in topography, morphological variations along the fibril, height profile or alignment between the native and labelled Tau fibrils.

**Figure 4.**
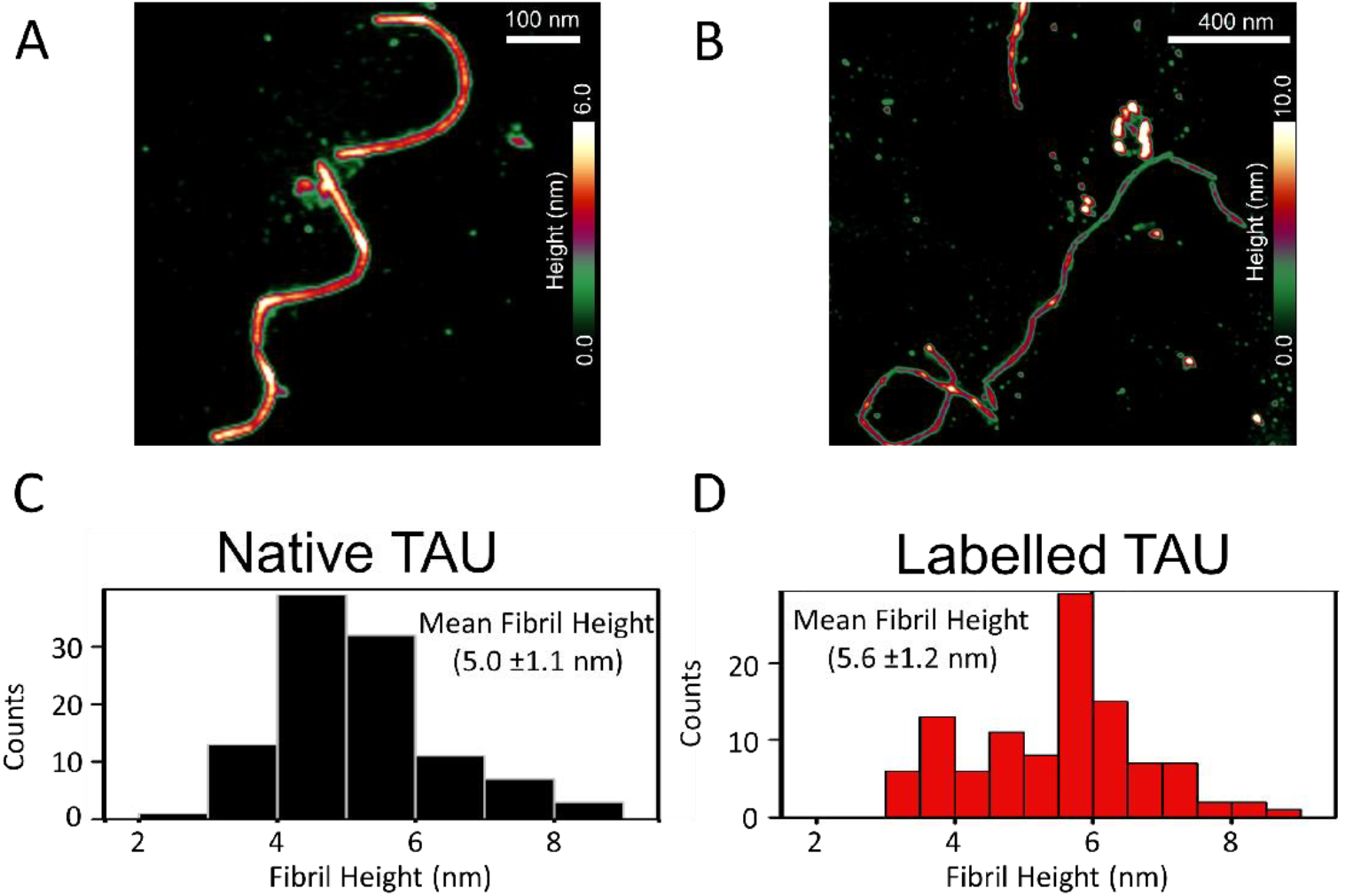
Atomic force microscopy (AFM) imaging and characterization of the height of Tau fibrils. **(A)** AFM image of native Tau fibrils. **(B)** AFM image showing the topography of labeled Tau fibrils. Height distribution (based on individual cross-sectional analyses of the AFM data obtained on native **(C)** and labelled **(D)** topographic images) of native Tau fibrils (black histogram) and labelled Tau fibrils (red histogram). A mean fibril height of 5.0±1.1 nm was obtained for the native Tau fibrils and a mean fibril height of 5.6 ± 1.2 nm was calculated for the labelled Tau fibrils from the height distribution plots.

## Conclusion

Using Sortase A-mediated labeling this work demonstrates that: 1) C-terminal labeling of Tau protein has minimal effect on secondary structure in solution, 2) both the labelled (i.e. Tau-647) and native-full length Tau protein exhibit similar aggregation kinetics, and 3) both the labelled and native Tau protein exhibit similar fibrillar morphology. We propose that the C-terminal labeling strategy of Tau protein presented here may therefore be useful for studies of Tau aggregation using (single-molecule) fluorescence methods with minimal effects on the structure of the native protein conformation.

## Acknowledgement

SA acknowledges financial support by the Swiss National Science Foundation’s “SPARK” funding (CRSK-3_195960). We acknowledge support from the National Institute on Aging of the National Institutes of Health under Award Numbers RF1AG062632 (JY) and RF1AG077802 (JY and MM). The content is solely the responsibility of the authors and does not necessarily represent the official views of the National Institutes of Health. MM acknowledges financial support from the Swiss National Science Foundation (Grant number: 200020_197239) and from the Adolphe Merkle Foundation, Switzerland.

## Materials and Methods

### Tau labeling

The pET30a plasmid containing GST-Tau-LPETGG-H6 was synthesized by GenScript. The protein was expressed by Oscar Vadas at the Protein and Peptides platform, University of Geneva, in E. Coli followed by purification. For labeling, 50 μM of purified Tau-LPETGG was mixed with 150 μM of the peptide GGGH_6_-Alexa647 (Bio-Synthesis Inc.) and 3 μM Sortase A5 (Active Motif) in 20 mM Tris, 150 mM NaCl, 10 mM CaCl_2_, 1 mM DTT pH 7.4. The mixture was incubated at 10°C for 3h and then loaded onto a His Spintrap column (GE Healthcare). After 1-hour incubation, the column was washed with 10 mM Tris, pH 7.8 100 mM NaCl, 10 mM imidazole followed by elution with 10 mM Tris, pH 7.8, 100 mM NaCl, 250 mM imidazole. The elute was concentrated using a vivaspin 500, 3000 MWCO centrifugal concentrator (Sartorius, Germany) and then purified on a Superdex200 increase 10/300 column (GE Healthcare) in 10 mM Tris, 100 mM NaCl, 1 mM DTT; pH 7.4 using a FPLC AKTA system (GE Healthcare). The fractions containing the purified labelled Tau protein were pooled, concentrated with a vivaspin 500, 3000 MWCO centrifugal concentrator, aliquoted, flash frozen in liquid N_2_ and stored at -80°C. At each step, samples were taken, mixed with SDS-PAGE loading buffer and loaded onto an SDS-polyacrylamide gel, for 40 min run at 150 V in order to verify the presence of the protein.

### ThT assays

The aggregation of native full-length Tau procured from Eurogentec and Tau-LPETGGGH_6_C-Alexa_647_ henceforth called Tau-A647 was monitored by measuring fluorescence spectra of ThT over time using a fluorometer from Horiba Scientific. For the Tau aggregation assay, a solution of 2 μM Tau was mixed with 0.5 μM heparin sodium salt (MW: 6’000-3’0000 g/mol), 1 μM of ThT in Tau aggregation buffer (25 mM sodium phosphate, 25 mM NaCl, 2.5 mM EDTA, 0.33 mM DTT, pH 6.8). The aggregation mix was incubated at 37°C, under constant agitation with 300 rpm in protein Lobind tubes (Eppendorf). Fluorescence emission spectra were measured at different time points (0h, 1h, 24h, 48h, 72h, 96h, 168h, 192h and 216h) by using an excitation wavelength of 412 nm and a slit of 5 nm in black quartz cuvettes (Hellma).

### CD spectroscopy

CD spectra were acquired on a Jasco J-810 CD spectropolarimeter using 0.1 cm quartz cuvettes (Hellma), operated at 20°C. To minimize buffer absorption, Tau samples were dialysed overnight against 10 mM sodium phosphate buffer, pH 7.8 in Slide-A-Lyzer MINI dialysis devices (100 ul). CD spectra were recorded from 190-260 nm at a scan speed of 20 nm/min and an increment of 1 nm. Four scans were recorded for each sample. The buffer was used as a blank for background subtraction.

### AFM

A total of 5 μl solution containing fibrils of native Tau or Tau-LPETGGGH_6_C-Alexa_647_ both at an equivalent monomer concentration of 100 nM in a buffer containing HEPES pH 7.4 was pipetted onto Si (111), n-type substrate. The sample was left to settle for 30 min followed by injecting 10 μl of pure water to rinse the buffer salt solution. Atomic force microscope images were acquired in tapping mode in pure water with a Bruker Multimode 8, E-scanner. The tip used was a Scanasyst-AIR, 0.4 N/m, 70 kHz (Bruker AFM probes). Images were recorded at a resolution of 1024 by 1024 pixels at a scan rate of 1 Hz. Analysis of AFM data was conducted using Bruker Nanoscope Analysis software.

### Electron Microscopy

Transmission electron microscopy images were recorded with a FEI Tecnai Spirit operating at 120 k. Carbon coated 300-mesh copper grids (Electron Microscopy Sciences, Hatfield) were plasma cleaned for 5 s using an oxygen plasma cleaner (Zepto RIE, Dienner), 5 μl of Tau protein sample in aggregation buffer (diluted 100 times) was pipetted on top of the grids and incubated for 2 min, the grids were then washed in a water droplet. Uranyl acetate (2% w/v) was added (3ul) and incubated for 2 min. Excess stain was blotted off with a filter paper and dried. We used ImageJ to estimate the width of Tau protein fibrils.^50^

## References

(1) Orr, M.E.; Sullivan A.C.; and Frost, B. A Brief Overview of Tauopathy: Causes, Consequences, and Therapeutic Strategies. Trends in Pharmacological Sciences 2017, 38(7): 637–648. DOI: https://doi.org/10.1016/j.tips.2017.03.011

(2) Lee, V.M.Y.; Goedert, M; and Trojanowski, J.Q. Neurodegenerative Tauopathies. Annual Reviews 2001, 24(1): 1121–1159. DOI: https://doi.org/10.1146/annurev.neuro.24.1.1121

(3) Spillantini, M.G; and Goedert, M. Tau protein pathology in neurodegenerative diseases. Trends in Neurosciences 1998, 21(10): 428–433. DOI: https://doi.org/10.1016/S0166-2236(98)01337-X

(4) Maccioni, R.B.; Muñoz, J.P.; and Barbeito, L. The Molecular Bases of Alzheimer’s Disease and Other Neurodegenerative Disorders. Archives of Medical Research 2001, 32(5): 367–381. DOI: https://doi.org/10.1016/S0188-4409(01)00316-2

(5) Cleveland, D.W.; Hwo, S.Y.; and Kirschner, M.W. Purification of tau, a microtubule-associated protein that induces assembly of microtubules from purified tubulin. J Mol Biol 1977, 116(2): 207–25. DOI: https://doi.org/10.1016/0022-2836(77)90213-3

(6) Drubin, D.G.; and Kirschner, M.W. Tau protein function in living cells. J Cell Biol, 1986, 103(6): 2739–46. DOI: https://doi.org/10.1083/jcb.103.6.2739

(7) Goedert, M. Tau filaments in neurodegenerative diseases. FEBS Lett 2018, 592(14): 2383–2391. DOI: 10.1002/1873-3468.13108.

(8) Verwilst, P.; Kim, H. S.; Kim, S.; Kang, C.; Kim, J. S. Shedding light on tau protein aggregation: the progress in developing highly selective fluorophores. Chem Soc Rev 2018, 47 (7), 2249–2265. DOI: 10.1039/c7cs00706j.

(9) Jeganathan, S.; von Bergen, M.; Brutlach, H.; Steinhoff, H. J.; Mandelkow, E. Global hairpin folding of tau in solution. Biochemistry 2006, 45 (7), 2283–2293. DOI: 10.1021/bi0521543.

(10) Luo, Y.; Ma, B.; Nussinov, R.; Wei, G. Structural Insight into Tau Protein’s Paradox of Intrinsically Disordered Behavior, Self-Acetylation Activity, and Aggregation. The Journal of Physical Chemistry Letters 2014, 5 (17), 3026–3031. DOI: 10.1021/jz501457f.

(11) Martin, L.; Latypova, X.; Terro, F. Post-translational modifications of tau protein: implications for Alzheimer’s disease. Neurochem Int 2011, 58 (4), 458–471. DOI: 10.1016/j.neuint.2010.12.023.

(12) Biernat, J.; Gustke, N.; Drewes, G.; Mandelkow, E.; Mandelkow, E. Phosphorylation of Ser262 strongly reduces binding of tau to microtubules: Distinction between PHF-like immunoreactivity and microtubule binding. Neuron 1993, 11 (1), 153–163. DOI: https://doi.org/10.1016/0896-6273(93)90279-Z.

(13) Noble, W.; Hanger, D. P.; Miller, C. C.; Lovestone, S. The importance of tau phosphorylation for neurodegenerative diseases. Front Neurol 2013, 4, 83. DOI: 10.3389/fneur.2013.00083.

(14) Tepper, K.; Biernat, J.; Kumar, S.; Wegmann, S.; Timm, T.; Hubschmann, S.; Redecke, L.; Mandelkow, E. M.; Muller, D. J.; Mandelkow, E. Oligomer formation of tau protein hyperphosphorylated in cells. J Biol Chem 2014, 289 (49), 34389–34407. DOI: 10.1074/jbc.M114.611368.

(15) Niewidok, B.; Igaev, M.; Sundermann, F.; Janning, D.; Bakota, L.; Brandt, R. Presence of a carboxy-terminal pseudorepeat and disease-like pseudohyperphosphorylation critically influence tau’s interaction with microtubules in axon-like processes. Mol Biol Cell 2016, 27 (22), 3537–3549. DOI: 10.1091/mbc.E16-06-0402.

(16) Lee, C. C.; Nayak, A.; Sethuraman, A.; Belfort, G.; McRae, G. J. A three-stage kinetic model of amyloid fibrillation. Biophys J 2007, 92 (10), 3448–3458. DOI: 10.1529/biophysj.106.098608.

(17) Wegmann, S.; Eftekharzadeh, B.; Tepper, K.; Zoltowska, K. M.; Bennett, R. E.; Dujardin, S.; Laskowski, P. R.; MacKenzie, D.; Kamath, T.; Commins, C.; et al. Tau protein liquid-liquid phase separation can initiate tau aggregation. EMBO J 2018, 37 (7). DOI: 10.15252/embj.201798049.

(18) Lasagna-Reeves, C. A.; Castillo-Carranza, D. L.; Sengupta, U.; Guerrero-Munoz, M. J.; Kiritoshi, T.; Neugebauer, V.; Jackson, G. R.; Kayed, R. Alzheimer brain-derived tau oligomers propagate pathology from endogenous tau. Sci Rep 2012, 2, 700. DOI: 10.1038/srep00700.

(19) Ghag, G.; Bhatt, N.; Cantu, D. V.; Guerrero-Munoz, M. J.; Ellsworth, A.; Sengupta, U.; Kayed, R. Soluble tau aggregates, not large fibrils, are the toxic species that display seeding and cross-seeding behavior. Protein Sci 2018, 27 (11), 1901–1909. DOI: 10.1002/pro.3499.

(20) Fitzpatrick, A. W. P.; Falcon, B.; He, S.; Murzin, A. G.; Murshudov, G.; Garringer, H. J.; Crowther, R. A.; Ghetti, B.; Goedert, M.; Scheres, S. H. W. Cryo-EM structures of tau filaments from Alzheimer’s disease. Nature 2017, 547 (7662), 185–190. DOI: 10.1038/nature23002.

(21) Wegmann, S.; Medalsy, I. D.; Mandelkow, E.; Müller, D. J. The fuzzy coat of pathological human Tau fibrils is a two-layered polyelectrolyte brush. Proc Natl Acad Sci 2013, 110 (4), E313–E321. DOI: 10.1073/pnas.1212100110.

(22) Fitzpatrick, A. W. P.; Falcon, B.; He, S.; Murzin, A. G.; Murshudov, G.; Garringer, H. J.; Crowther, R. A.; Ghetti, B.; Goedert, M.; Scheres, S. H. W. Cryo-EM structures of tau filaments from Alzheimer’s disease. Nature 2017, 547 (7662), 185–190. DOI: 10.1038/nature23002.

(23) von Bergen, M.; Friedhoff, P.; Biernat, J.; Heberle, J.; Mandelkow, E. M.; Mandelkow, E. Assembly of tau protein into Alzheimer paired helical filaments depends on a local sequence motif ((306)VQIVYK(311)) forming beta structure. Proc Natl Acad Sci 2000, 97 (10), 5129–5134. DOI: 10.1073/pnas.97.10.5129.

(24) Ganguly, P.; Do, T. D.; Larini, L.; LaPointe, N. E.; Sercel, A. J.; Shade, M. F.; Feinstein, S. C.; Bowers, M. T.; Shea, J.-E. Tau Assembly: The Dominant Role of PHF6 (VQIVYK) in Microtubule Binding Region Repeat R3. The Journal of Physical Chemistry B 2015, 119 (13), 4582–4593. DOI: 10.1021/acs.jpcb.5b00175.

(25) Bergen, M. v.; Friedhoff, P.; Biernat, J.; Heberle, J.; Mandelkow, E.-M.; Mandelkow, E. Assembly of tau protein into Alzheimer paired helical filaments depends on a local sequence motif ((306)VQIVYK(311)) forming beta structure. 2000, 97 (10), 5129–5134. DOI: doi:10.1073/pnas.97.10.5129.

(26) Friedhoff, P.; von Bergen, M.; Mandelkow, E. M.; Davies, P.; Mandelkow, E. A nucleated assembly mechanism of Alzheimer paired helical filaments. Proc Natl Acad Sci 1998, 95 (26), 15712–15717. DOI: 10.1073/pnas.95.26.15712.

(27) Schweers, O.; Mandelkow, E. M.; Biernat, J.; Mandelkow, E. Oxidation of cysteine-322 in the repeat domain of microtubule-associated protein tau controls the in vitro assembly of paired helical filaments. Proc. Natl. Acad. Sci. U. S. A. 1995, 92 (18), 8463–8467.

(28) Prifti, E.; Tsakiri, E. N.; Vourkou, E.; Stamatakis, G.; Samiotaki, M.; Papanikolopoulou, K. The Two Cysteines of Tau Protein Are Functionally Distinct and Contribute Differentially to Its Pathogenicity in Vivo. The Journal of Neuroscience. 2021, 41 (4), 797–810. DOI: 10.1523/JNEUROSCI.1920-20.202028.

(29) Karikari, T. K.; Nagel, D. A.; Grainger, A.; Clarke-Bland, C.; Hill, E. J.; Moffat, K. G. Preparation of stable tau oligomers for cellular and biochemical studies. Anal Biochem 2019, 566, 67–74. DOI: 10.1016/j.ab.2018.10.013.

(30) Di Primio, C.; Quercioli, V.; Siano, G.; Rovere, M.; Kovacech, B.; Novak, M.; Cattaneo, A. The Distance between N and C Termini of Tau and of FTDP-17 Mutants Is Modulated by Microtubule Interactions in Living Cells. Front Mol Neurosci 2017, 10, 210. DOI: 10.3389/fnmol.2017.00210.

(31) Michel, C. H.; Kumar, S.; Pinotsi, D.; Tunnacliffe, A.; St George-Hyslop, P.; Mandelkow, E.; Mandelkow, E. M.; Kaminski, C. F.; Kaminski Schierle, G. S. Extracellular monomeric tau protein is sufficient to initiate the spread of tau protein pathology. J Biol Chem 2014, 289 (2), 956–967. DOI: 10.1074/jbc.M113.515445.

(32) Hinrichs, M. H.; Jalal, A.; Brenner, B.; Mandelkow, E.; Kumar, S.; Scholz, T. Tau Protein Diffuses along the Microtubule Lattice*. Journal of Biological Chemistry 2012, 287 (46), 38559–38568. DOI: https://doi.org/10.1074/jbc.M112.369785.

(33) Elbaum-Garfinkle, S.; Ramlall, T.; Rhoades, E. The Role of the Lipid Bilayer in Tau Aggregation. Biophysical Journal 2010, 98 (11), 2722–2730. DOI: https://doi.org/10.1016/j.bpj.2010.03.013.

(34) Jeganathan, S.; von Bergen, M.; Mandelkow, E.-M.; Mandelkow, E. The Natively Unfolded Character of Tau and Its Aggregation to Alzheimer-like Paired Helical Filaments. Biochemistry 2008, 47 (40), 10526–10539. DOI: 10.1021/bi800783d.

(35) Ramachandran, G.; Udgaonkar, J. B. Understanding the kinetic roles of the inducer heparin and of rod-like protofibrils during amyloid fibril formation by Tau protein. The Journal of biological chemistry 2011, 286 (45), 38948–38959. DOI: 10.1074/jbc.M111.271874

(36) Goedert, M.; Jakes, R.; Spillantini, M. G.; Hasegawa, M.; Smith, M. J.; Crowther, R. A. Assembly of microtubule-associated protein tau into Alzheimer-like filaments induced by sulphated glycosaminoglycans. Nature 1996, 383 (6600), 550–553. DOI: 10.1038/383550a0.

(37) Perez, M.; Valpuesta, J. M.; Medina, M.; Montejo de Garcini, E.; Avila, J. Polymerization of tau into filaments in the presence of heparin: the minimal sequence required for tau-tau interaction. J Neurochem 1996, 67 (3), 1183–1190. DOI: 10.1046/j.1471-4159.1996.67031183.x.

(38) Kampers, T.; Friedhoff, P.; Biernat, J.; Mandelkow, E. M.; Mandelkow, E. RNA stimulates aggregation of microtubule-associated protein tau into Alzheimer-like paired helical filaments. FEBS Lett 1996, 399 (3), 344–349. DOI: 10.1016/s0014-5793(96)01386-5.

(39) Friedhoff, P.; Schneider, A.; Mandelkow, E. M.; Mandelkow, E. Rapid assembly of Alzheimer-like paired helical filaments from microtubule-associated protein tau monitored by fluorescence in solution. Biochemistry 1998, 37 (28), 10223–10230. DOI: 10.1021/bi980537d.

(40) Witte, M. D.; Wu, T.; Guimaraes, C. P.; Theile, C. S.; Blom, A. E.; Ingram, J. R.; Li, Z.; Kundrat, L.; Goldberg, S. D.; Ploegh, H. L. Site-specific protein modification using immobilized sortase in batch and continuous-flow systems. Nat Protoc 2015, 10 (3), 508–516. DOI: 10.1038/nprot.2015.026.

(41) Dillard, K. E.; Schaub, J. M.; Brown, M. W.; Saifuddin, F. A.; Xiao, Y.; Hernandez, E.; Dahlhauser, S. D.; Anslyn, E. V.; Ke, A.; Finkelstein, I. J. Sortase-mediated fluorescent labeling of CRISPR complexes. Methods Enzymol 2019, 616, 43–59. DOI: 10.1016/bs.mie.2018.10.031.

(42) Theile, C. S.; Witte, M. D.; Blom, A. E.; Kundrat, L.; Ploegh, H. L.; Guimaraes, C. P. Site-specific N-terminal labeling of proteins using sortase-mediated reactions. Nat Protoc 2013, 8 (9), 1800–1807. DOI: 10.1038/nprot.2013.102.

(43) Xue, C.; Lin, T. Y.; Chang, D.; Guo, Z. Thioflavin T as an amyloid dye: fibril quantification, optimal concentration and effect on aggregation. 2017, 4 (1), 160696. DOI: doi:10.1098/rsos.160696.

(44) Li, J.; Zhang, Y.; Soubias, O.; Khago, D.; Chao, F. A.; Li, Y.; Shaw, K.; Byrd, R. A. Optimization of sortase A ligation for flexible engineering of complex protein systems. J Biol Chem 2020, 295 (9), 2664–2675. DOI: 10.1074/jbc.RA119.012039.

(45) Chen, I.; Dorr, B. M.; Liu, D. R. A general strategy for the evolution of bond-forming enzymes using yeast display. Proc Natl Acad Sci U S A 2011, 108 (28), 11399–11404. DOI: 10.1073/pnas.1101046108.

(46) Wiedemann, C.; Bellstedt, P.; Görlach, M. CAPITO--a web server-based analysis and plotting tool for circular dichroism data. Bioinformatics (Oxford, England) 2013, 29 (14), 1750–1757. DOI: 10.1093/bioinformatics/btt278

(47) Gade Malmos, K.; Blancas-Mejia, L. M.; Weber, B.; Buchner, J.; Ramirez-Alvarado, M.; Naiki, H.; Otzen, D. ThT 101: a primer on the use of thioflavin T to investigate amyloid formation. Amyloid 2017, 24 (1), 1–16. DOI: 10.1080/13506129.2017.1304905

(48) Morozova, O. A.; March, Z. M.; Robinson, A. S.; Colby, D. W. Conformational features of tau fibrils from Alzheimer’s disease brain are faithfully propagated by unmodified recombinant protein. Biochemistry 2013, 52 (40), 6960–6967. DOI: 10.1021/bi400866w.

(49) Zhang, W.; Falcon, B.; Murzin, A. G.; Fan, J.; Crowther, R. A.; Goedert, M.; Scheres, S. H. Heparin-induced tau filaments are polymorphic and differ from those in Alzheimer’s and Pick’s diseases. Elife 2019, 8. DOI: 10.7554/eLife.43584.

(50) Schneider, C. A.; Rasband, W. S.; Eliceiri, K. W. NIH Image to ImageJ: 25 years of image analysis. Nature Methods 2012, 9 (7), 671–675. DOI: 10.1038/nmeth.2089.

